# MxB N-Terminus Adopts a Stable *α*-Helix to Engage the HIV-1 Capsid Trimer Interface

**DOI:** 10.64898/2026.09.03.749210

**Authors:** Yanan Zhu, Juan S. Rey, Juan Shen, Juan R. Perilla, Peijun Zhang

## Abstract

The HIV-1 capsid, composed of capsid (CA) proteins arranged into a conical surface lattice, serves as a critical platform for host–pathogen interactions. Many host cofactors regulate HIV-1 infection by selectively recognizing the higher-order CA lattice rather than individual monomers or capsomers. Among these, Myxovirus resistance protein B (MxB) is a key cellular restriction factor that binds the capsid via its N-terminal fragment containing an arginine-rich motif (RRR), blocking HIV-1 infection at early infection stages. However, the structural basis of this interaction has remained elusive. Here, we present a structural characterization of the CA lattice in complex with the MxB N-terminal fragment using cryo-electron tomography and subtomogram averaging. Combining cryoEM structures with EM-guided all-atom molecular dynamics simulations, we identify residue-specific interactions between MxB and the CA trimer interfaces. Notably, residues 10-20 of the MxB N-terminus adopt an α-helical conformation that stabilizes binding at the CA trimer interface, revealing a previously unrecognized mode of capsid engagement. These findings provide new insight into arginine-rich motif–mediated recognition by MxB and other host factors and establish a structural framework for the development of capsid-targeting antiviral therapeutics.

## Introduction

Human immunodeficiency virus type 1 (HIV-1) is a retrovirus that causes acquired immune deficiency syndrome (AIDS) by depleting CD4^+^ T cells and compromising host immunity (*1*). The mature HIV-1 virion contains a conical capsid composed of around 200 hexameric and 12 pentameric capsid protein (CA) subunits (*2–5*). The viral genome is housed within the capsid, which protects it from host immune defences during the early stages of the viral replication cycle (*4*). The stability and disassembly of the capsid are tightly regulated by interactions with host cell factors(*6–10*), including inositol hexakisphosphate (IP6) (*11–14*), cyclophilin A (CypA) (*15–17*), tripartite motif-containing protein 5α (Trim5α) (*10, 18, 19*), TrimCyp (*20, 21*), Myxovirus resistance protein B (MxB) (*22–25*) and cleavage and polyadenylation specificity factor 6 (CPSF6) (*7, 26–29*). Many of these factors recognize high-order structural features of the capsid lattice rather than individual CA capsomers, underscoring the importance of capsid geometry in viral replication and immune evasion (*6, 30*).

Among the capsid-interacting host factors, CypA and Trim5α have been extensively characterized for their ability to bind the HIV-1 capsid and modulate infection. CypA binds to exposed loops of CA lattice, stabilizing curvature-dependent interfaces that influence capsid stability and nuclear entry (*15, 17*). Trim5α assembles into hexagonal lattices on the surface of the capsid through its C-terminal SPRY domain, creating a physical barrier that affects viral uncoating (*31*). Host restriction factor MxB is a dynamin-like guanosine triphosphatase (GTPase) and is part of a group of interferon-stimulated genes (ISG) produced by the cell to combat viral infection. This group includes the well-known ISG myxovirus resistance protein A (MxA) (*32, 33*). Unlike its paralog MxA, which restricts influenza virus, thogotovirus, and vesicular stomatitis virus by targeting viral nucleocapsids or replication complexes in the cytoplasm (*33*), MxB preferentially restricts HIV-1, hepatitis C virus (HCV) and herpesviruses by acting at the level of the viral capsid and nuclear import, particularly in the case of HIV-1 (*23, 25, 34–39*). MxA and MxB share 63% sequence identity and are structurally similar, each comprising an N-terminal domain (NTD), a GTPase domain, and a C-terminal stalk domain that mediates dimerization (*24, 25, 40, 41*). The functional divergence between MxA and MxB is attributed to the unique N-terminal extension of MxB (*35, 42, 43*), which is absent in MxA. This region contains a conserved triple-arginine motif (RRR_11_-_13_) that directly engages inter-hexamer interfaces in the capsid lattice (*22, 24, 25, 35*). However, the structural basis of this interaction remains unresolved, limiting our understanding of how the MxB N-terminus selectively recognizes the HIV-1 capsid and mediates viral restriction. Elucidating this mechanism is crucial for defining the molecular determinants of MxB’s antiviral activity and for guiding the development of therapeutics that mimic or enhance this natural defense.

In this study, we employed cryo-electron tomography (cryoET) and subtomogram averaging (STA) to investigate the molecular interactions between MxB and the HIV-1 capsid. CryoET STA structures of MxB_1-35_ N-terminus in complex with the mature capsid lattice revealed distinct MxB densities located at CA trimer interfaces. All-atom molecular dynamics simulations of the MxB**_1-35_**-CA trimer complex identified key interactions between positively charged residues in MxB and glutamate and threonine residues in CA at the trimer interface. These interactions stabilize residues 10–20 of MxB in an α-helical conformation at the binding site. Together, these findings define the structural basis of the MxB-capsid interaction and provide mechanistic insight into how this restriction factor interferes with viral infection. More broadly, they advance our understanding of capsid-targeting host defences and may inform the development of antiviral strategies that exploit MxB’s unique mode of binding.

## Results

### CryoET STA study of the mature CA hexamer from CA-NC tubular assemblies

CA-NC tubular assemblies, which contain a mature CA lattice, are a well-established model for investigating HIV-1 capsid-cofactor interactions (*3, 15, 17*). This system avoids the use of high salt (∼1 M NaCl), required for CA tube assembly, or high concentrations of IP6 (>1 mM), used to stabilize capsid-like particles (CLPs). Such non-physiological conditions can influence host cofactor binding to the capsid. Despite their utility, a high-resolution structure of mature CA-NC tubular assemblies has not previously been reported. Using cryoET and STA, we determined the density map of the CA hexamer from CA-NC tubes at a global resolution of 4.9Å (Fig. 1, Fig. S1a, Fig. S2a and Table S1). The CA density is well resolved, whereas the NC density is poorly defined, consistent with its intrinsic flexibility, a property essential for genomic RNA packaging during viral assembly (*44*).

**Figure 1.**
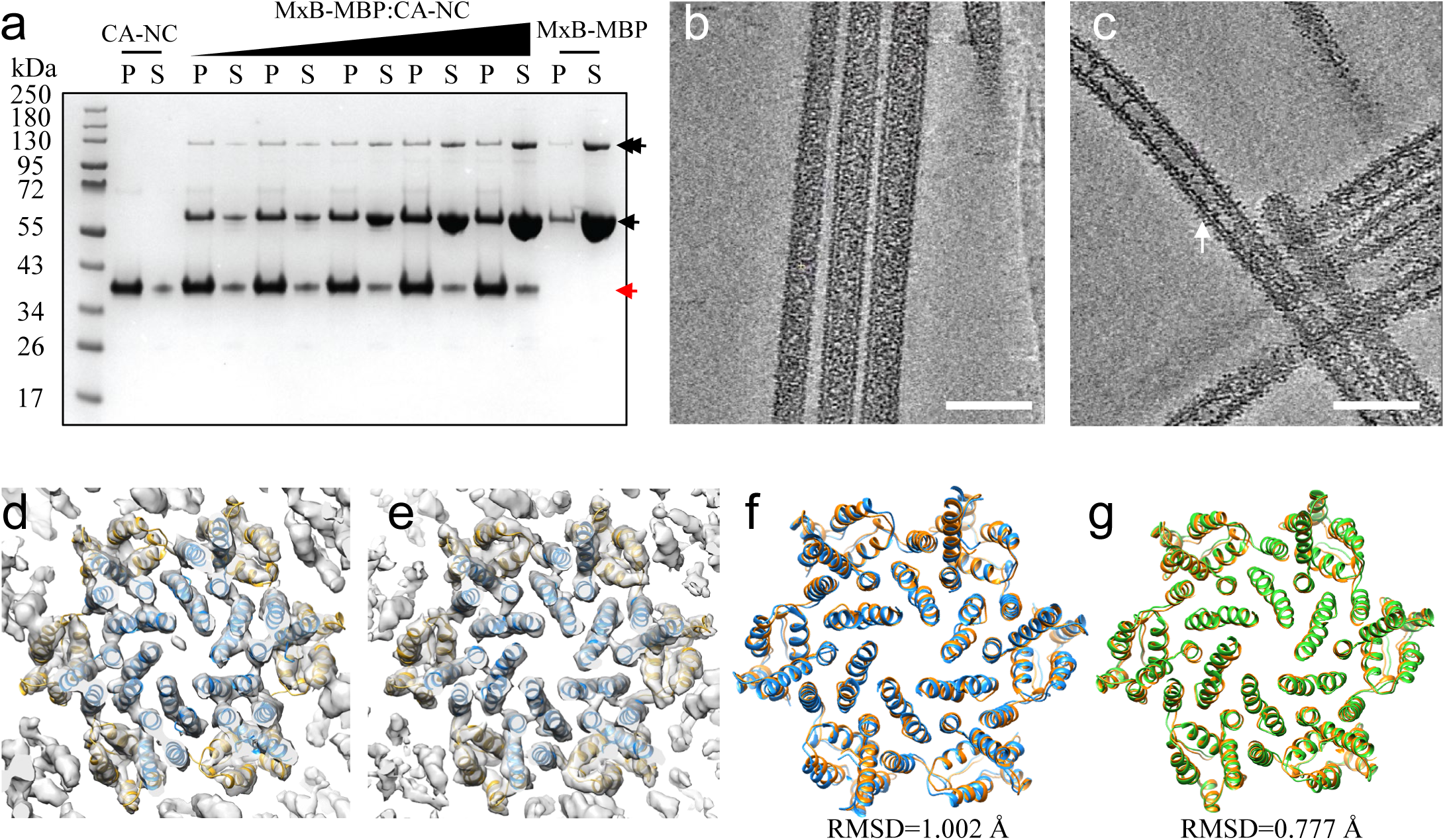
Structures of mature CA hexamer and in complex with MxB_1-35_. (a) SDS PAGE analysis of Sup/pellet assay for the binding of MBP-tagged MxB_1-35_ to preassembled CA-NC tubes at different molar ratios (from left to right, CA-NC only, MxB_1-35_ :CA-NC 0.125, 0.25, 0.5, 1 and 2, MxB_1-35_ only respectively). Single and double black arrows mark MxB_1-35_ -MBP and MxB_N_-MBP dimer, respectively; red arrow marks CA-NC. (b-c) Tomographic slices of CA-NC tube in the absence (b) and presence (c) of MxB_1-35_ at molar ratio 1:1. Scale bars, 100 nm. The arrow points additional densities. (d-e) CryoET STA structures of CA hexamers from CA-NC tubes without (d) and with (e) MxB_1-35_ binding superimposed with CA atomic model (PDB:4XFX) with CA_NTD_ (blue) and CA_CTD_ (orange) fitted separately. (f) Pairwise comparison of CA hexamer structures derived from CA-NC (orange) and CA (blue) tubes. (g) Pairwise comparison of CA hexamer structures derived from CA-NC (orange) and CA-NC/MxB_1-35_ tubes (green). The backbone RMSD values are indicated.

An atomic model of the mature CA hexamer lattice was generated by EM density-guided rigid-body fitting in coot, followed by real-space refinement in Phenix. The CA hexamer derived from CA–NC tubes closely resembles structures of canonical CA tubes assembled under high-salt conditions (*17*) (Cα RMSD = 1.0 Å, Fig. 1f), CLPs stabilized with high IP6 concentrations (RMSD range 0.93 to 1.016 Å) (*45, 46*), and the native mature capsid (*13*). These structural comparisons validate CA-NC tubular assemblies as a physiologically relevant platform for studying capsid–cofactor interactions.

### MxB_1-35_ binds to the CA-NC tubes

Because full-length MxB exhibits poor biochemical behavior, we used a capsid-binding construct comprising the N-terminal 35 residues of MxB fused to maltose-binding protein (MBP) and a GCN4 dimerization domain to mimic the native MxB dimer, referred to hereafter as MxB_1-35_ (*22*). Co-pelleting assays with CA–NC tubes confirmed that MxB_1-35_ binds directly to CA–NC assemblies (Fig. 1a), consistent with previous report (*22*). Binding saturated at a molar ratio of approximately 0.25; therefore, complexes formed at this ratio were used for cryo-ET analysis. Tomographic slices from reconstructed cryoET volumes revealed additional densities decorating the tube surface (Fig.1c) compared to CA-NC tubes alone (Fig. 1b), indicating that these densities correspond to the bound MxB1-35 fusion protein.

We performed cryoET STA of CA hexamers within MxB_1-35_-bound CA-NC tubes. A low-pass-filtered 30 Å map of a 7-CA hexamer assembly was used as a template for particle picking and initial alignment (Fig. S1b). Subsequent alignment and refinement yielded a CA hexamer structure at a global resolution of 4.4 Å from 129,046 particles (Fig. 1e, Fig. S2a and Table S1). Densities corresponding to both the CA N-terminal domain (CA_NTD_) and C-terminal domain (CA_CTD_) were well resolved (Fig. 1e and Fig. S2b-c), whereas NC density remained poorly defined, consistent with its intrinsic flexibility.

To further improve resolution, we performed cryoEM single-particle analysis (SPA) of CA hexamers from MxB_1-35_-bound CA-NC tubes (Fig. S3-4). The resulting map at 2.8 Å resolution shows well-resolved side-chain densities, enabling accurate atomic model building (Fig. S3-4 and Table S2). The SPA-derived CA hexamer structure closely matches the cryoET/STA model (Cα RMSD = 0.85 Å, Fig. S4b). Comparison of CA hexamer models in the presence and absence of MxB_1-35_ revealed only minor structural differences (Cα RMSD = 0.78 Å; Fig. 1g), indicating that MxB binding does not substantially perturb the CA hexamer structure or overall lattice organization.

### Dimeric MxB_1-35_ binds pairwise to and remodels the capsid trimer interface

Previous studies have implicated the trimer interface as the MxB binding site (*22*), we next focused our structural analysis on this region. Extensive focused alignment and classification of the trimer interface, as well as density subtraction using a cryoEM SPA approach, did not yield clearly distinguishable densities. However, the cryoET/STA map revealed a distinct extra density bridging two adjacent H10 helices at the trimer interface (Fig. S5a). This density was absent in the STA map of CA-NC tubes lacking MxB_1-35_ (Fig. S5b), providing evidence for MxB engagement at this site.

To improve local resolution and better resolve this interaction, we re-centred subvolumes on the trimer interface and performed focused refinement using an alignment mask encompassing this region, followed by 3D classification (Fig. S1b). Two major classes were identified. Class 1 (5.3 Å resolution), representing 16.6% of particles (Fig. S1b, S2a,d and Table S1), displayed additional density extending from the CA dimer interface (near E187) across the H10 helices to the trimer interface (near E213 and P207) (Fig. 2a-b, red density). Class 2 (4.5 Å resolution), comprising 83.4% of particles, lacked this extra density (Fig. S1b, S2a,e). This heterogeneity suggests partial occupancy of MxB at trimer interfaces. Although MxB binding does not substantially alter the overall CA hexamer structure (Fig. 1g), localized conformational changes were observed at the trimer interface. Notably, the CA_CTD_ helices of one subunit exhibited pronounced displacement, with an RMSD of 4.95 Å (Fig. 2c), indicating a structural response to MxB engagement.

**Figure 2.**
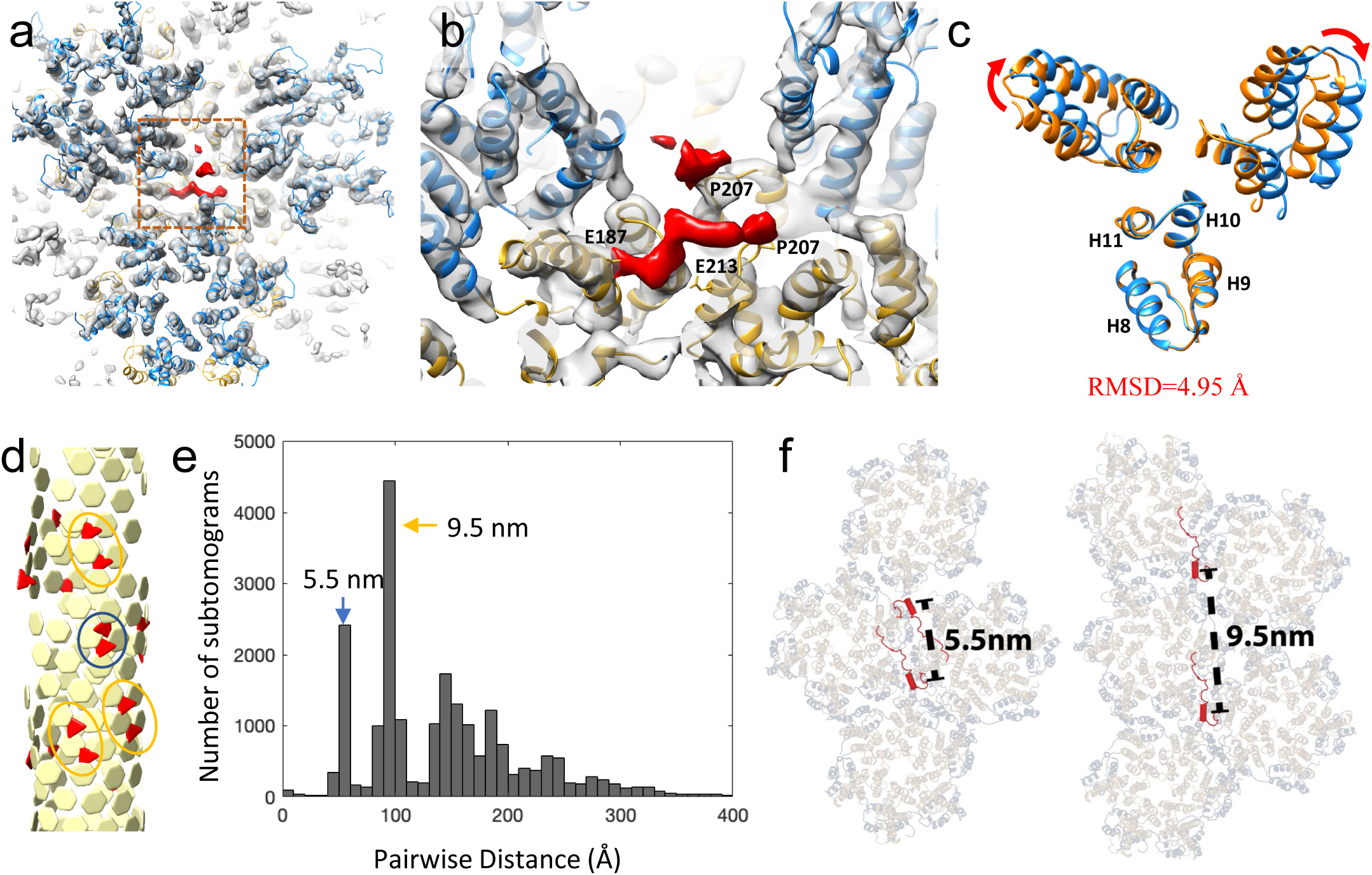
CryoET STA Structure of MxB_1-35_ and CA-NC tube complexes. (a) MxB_1-35_ density, segmented in red, above the trimer interface. (b) A close-up view of the MxB_1-35_ density, with nearby CA trimer interface residues labeled (E187, E213, P207). (c) A comparison of CA trimer interfaces in the absence (orange) and presence (blue) of MxB_1-35_. Structures are aligned to one CA-CTD (the lower one). (d) A geometrical model of CA hexamer positions (yellow) and MxB_1-35_ positions (red) mapped onto a CA-NC tube through cryoET STA analysis. Distinct pairs of MxB_1-35_ densities are circled in blue for short and in orange for intermediate distances. (e) Pairwise distance distribution of MxB_1-35_ positions from all subtomograms. The two most frequent distances are 5.5 nm (blue arrow) and 9.5 nm (orange arrow). (f) Models of MxB_1-35_ binding on a CA hexamer lattice based on the predominant observed distances between MxB_1-35_ -occupied trimer interfaces.

Given that native MxB functions as a dimer (*22, 25*), and our construct MxB_1-35_-MBPdi is designed to mimic this dimeric state, we next examined the spatial distribution of binding on CA-NC tubes. Mapping the Class 1 particle positions back onto the original tomograms revealed that MxB-occupied trimer interfaces frequently occurred in pairs (Fig. 2d). Nearest-neighbour distance analysis of MxB-occupied trimer centers showed two prominent peaks: the first at 5.5 nm, corresponding to the distance between adjacent trimers (Fig. 2d-f, blue circle and arrow), and the second at 9.5 nm, matching the centre-to-centre distance between neighbouring hexamers (Fig. 2d-f, orange ovals and arrow). These observations suggest that two adjacent trimer sites can be simultaneously engaged by a single MxB dimer (Fig. 2f).

### MxB binding is driven by electrostatic interactions centred on the RRR_11-13_ motif

Guided by cryoET STA density, we performed molecular dynamics (MD) simulations to characterize the behaviour of the MxB N-terminal region at the CA trimer interface. The 35-residue MxB N-terminus was initially generated de-novo using AlphaFold2 as a template and subsequently modelled into the density observed on CA-NC tubes in complex with MxB_1-35_ using RosettaCM density-guided comparative modelling (see Methods; Fig. S6a-e). The resulting model, comprising MxB bound to a CA trimer of dimers, was solvated and ionized under physiologically relevant conditions and subjected to MD simulations totalling 1.6 μs of sampling time (Fig. 3a, Fig. S6f). Throughout the simulation, we calculated the root-mean-square fluctuation (RMSF) of the Cα atoms for each residue in MxB_1-35_ (Fig. 3b). Residues Met1 to Tyr21 exhibited low RMSF values (<2 Å), indicating structural stability, whereas residues beyond this region were substantially more flexible. Notably, Arg11 to Ser18 consistently adopted an α-helical conformation, with Arg19 and Lys20 partially maintaining helicity while intermittently transitioning into coil and turn conformations (Fig. 3c). These findings are consistent with the cryoET density at the CA trimer interface (Fig. 2), which reveals a short helix-like feature while the remainder of the MxB N-terminus remains unresolved.

**Figure 3.**
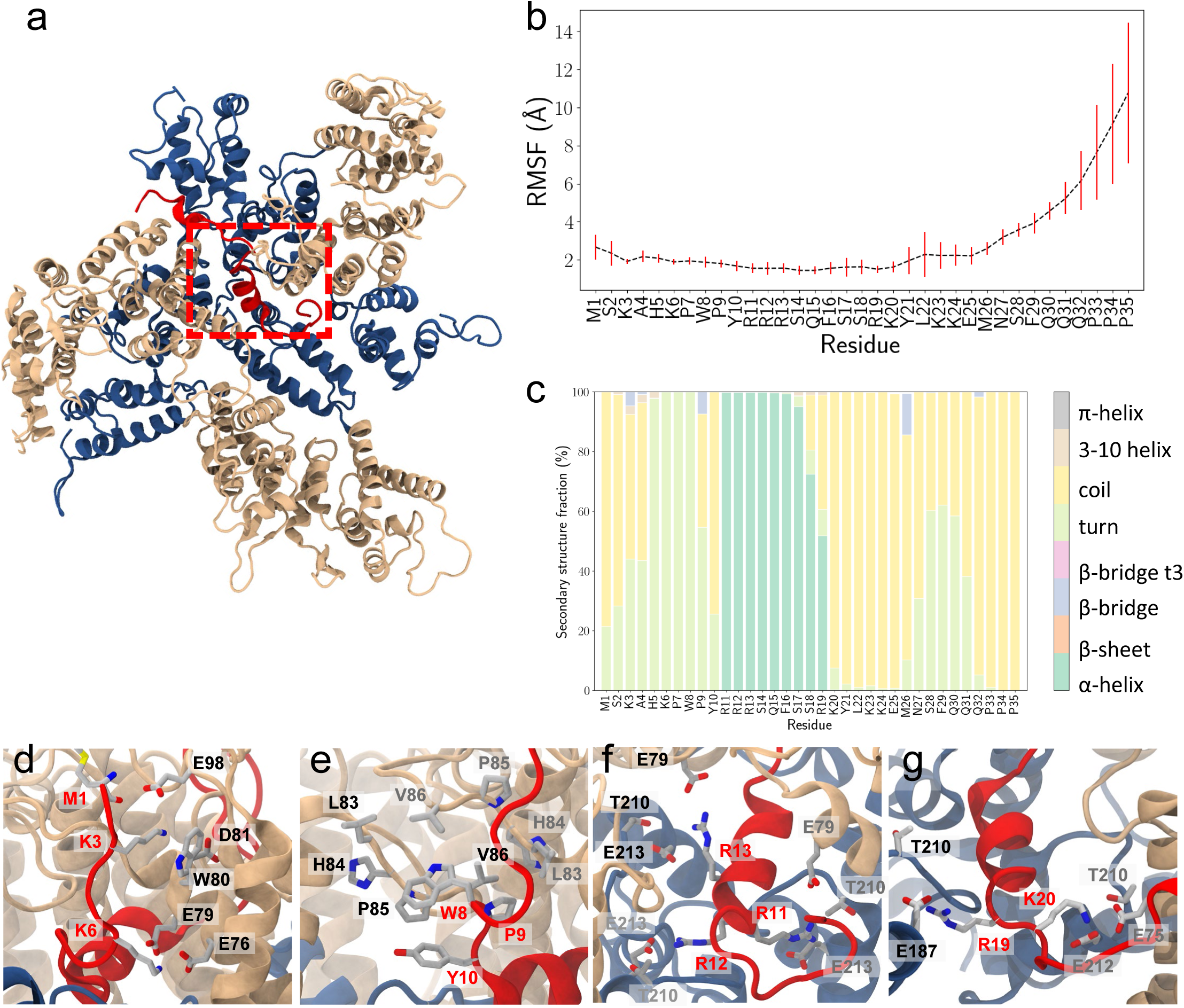
MxB_1-35_ interactions with the CA trimer interface from all-atom molecular dynamics (MD) simulations. (a) Representative conformation of MxB_1-35-_CA trimer of dimers complex from MD simulations. A dashed red box highlights the MxB binding pocket at the CA trimer interface. CA-NTD is colored in gold, CA-CTD in dark blue, and MxB_1-35_ in red. (b) Root-mean square-fluctuation (RMSF) trace for the displacement of Cα atoms in MxB_1-35_ over the course of the simulation. (c) Secondary structure fraction of MxB_1-35_ residues throughout the MD simulations. Residues R11 to S18 predominantly adopt an α-helix conformation. (d-g) Interactions between MxB_1-35_ and CA that stabilize the MxB_11-20_ α-helix within the trimer interface pocket:. (d) salt-bridge interactions involving the MxB N-terminal M1, K3 and K6; (e) hydrophobic contacts involving W8, P9 and Y10; (f) salt bridge interactions involving the RRR_11-13_ motif; and (g) salt bridge interactions involving R19 and K20 at the end of the α-helix. MxB_1-35_ and CA residues are shown as sticks, colored by atomic element and labeled in red and black, respectively. When multiple CA chains are involved in the interactions, residues from additional CA chains are labeled in gray.

Although the STA density does not provide sufficient resolution to directly resolve residue side chains or unambiguously identify MxB-CA residue interactions, we analyzed the MD simulation trajectories to characterize the network of contacts that stabilize MxB_1-35_ at the trimer interface in our model. To this end, we quantified occupancy of contacts between MxB_1-35_ and CA residues throughout the MD trajectory (Fig. 3d-g; Table S3). High contact occupancies were observed between MxB Lys3 and CA Glu98 (95%), as well as between MxB Lys6 and CA residues Glu76 (98%) and Glu79 (99%), consistent with favourable electrostatic interactions between the positively charged lysine residues in MxB_1-35_ with negatively charged residues in CA. MxB1-35 Lys3 was also frequently positioned adjacent to CA Trp80 (99% occupancy), supporting a cation–π interaction between these residues. In addition, the positively charged N-terminal amino group of Met1 transiently engaged Glu98 (66%), contributing to the stabilization of the MxB_1-35_ N-terminal region (Fig. 3d). Frequent contacts between hydrophobic residues were also observed. MxB Trp8 and Pro9 residues remained in close proximity to CA residues Leu83, Pro85, and Val86 at the base of the cyclophilin A (CypA) binding loop, with occupancies of 92.6%, 54.7%, and 81.6%, respectively, supporting the formation of a hydrophobic interface (Fig. 3e).

Immediately downstream, according to our model, the highly conserved and positively charged RRR_11_-_13_ motif formed contacts with negatively charged residue Glu213 from all three CA monomers at the trimer interface (Arg11-Glu213: 54.2%, Arg12-Glu213: 84.1%, Arg13-Glu213: 61.2%) and transiently interacted with Thr210 (Arg11-Thr210: 71.8%, Arg12-Thr210: 28.1%, Arg13-Thr210: 27.8%) (Fig. 3f). During the simulations, the RRR_11_-_13_ motif consistently formed the first turn of a stable α-helix. This structural element is critical for MxB-capsid binding and complex stability, consistent with prior reports demonstrating that mutation of the RRR_11_-_13_ to AAA severely impairs MxB binding and antiviral activity (*22, 47*). In our model, the α-helix extends for approximately three helical turns and is capped by Arg19 and Lys20. Residue Arg19 transiently formed contacts with the CA Gly208 backbone (72.8%) and Thr210 (55.0%), whereas Lys20 maintained high-occupancy contacts with Glu75 (72.3%) and transiently contacted Glu 212 (46.5%). Together, these interactions are consistent with a network of electrostatic and hydrogen-bonding interactions that stabilizes the α-helical segment within the trimer interface (Fig. 3g). The stabilizing interactions are supported by the low RMSF values observed across the helical region (Fig. 3b) and by the corresponding density in the cryoET map (Fig. 2b).

To further assess the relative importance of individual MxB_1-35_ and CA residues, we performed MM/GBSA calculations to estimate the per-residue energetic contributions to MxB_1-35_ binding (Fig. S7). Among MxB_1-35_ residues, positively charged residues provided the highest energetic contributions to binding (Fig. S7a), with the RRR_11_-_13_ motif exhibiting the largest estimated energetic contributions (Arg11: −6.3 kcal/mol , Arg12: −10.6 kcal/mol, Arg13: −4.6 kcal/mol), followed by Lys6 (−9.6 kcal/mol), Lys3 (−5.6 kcal/mol), Arg19 (−5.0 kcal/mol), and Lys20 (−4.8 kcal/mol). Hydrophobic and aromatic residues also made favourable energetic contributions, including Trp8 (−3.1 kcal/mol), Pro9 (−1.9 kcal/mol), and Tyr10 (−5.0 kcal/mol).

The energetic contributions of CA residues varied among the three monomers because the orientation of the bound MxB_1-35_ segment breaks the threefold symmetry of the trimer interface (Fig. S7b). Among CA residues, Glu213 consistently exhibited favourable energetic contributions to binding across all three CA chains (mean ΔG = −2.8 ± 1.8 kcal/mol), in agreement with its high occupancy contacts with the RRR_11-13_ motif. Other glutamate residues at the trimer interface also contributed favourably to binding, including Glu75 (−0.3 kcal/mol), Glu76 (−0.4 kcal/mol), Glu98 (−1.2 kcal/mol), and Glu212 (−0.6 kcal/mol). In contrast, Glu79 and Glu187 exhibited slightly unfavourable average energetic contributions (0.1 and 0.6 kcal/mol, respectively). Finally, the hydrophobic residues Trp80, Leu83, Pro85 and Val86 also contributed favourably to binding (−1.6, −2.0, −1.2, and −1.4 kcal/mol, respectively), further supporting the hydrophobic contacts identified throughout the MD trajectories.

To place these findings in structural context, we compared our MxB_1-35_ bound with curved-lattice assembly with previously reported simulations of MxB_1-35_ with a flat lattice lacking experimental constraints (Fig.4a) (*22*). In the curved CA lattice, the trimer interface forms a more open binding pocket that accommodates the full three-turn α-helix of MxB_1-35_, stabilized by electrostatic interactions between basic MxB residues and acidic CA patches. In contrast, the flatter lattice exhibits more tightly packed CA helices at the trimer interface, displacing MxB_1-35_ downward (Fig. 4a). In these simulations, MxB_1-35_ forms only a single helical turn encompassing the RRR_11_-_13_ motif, likely due to the spatial constraints imposed by the flatter geometry (Fig. 4a). These comparisons suggest that capsid curvature is a critical determinant of MxB engagement.

**Figure 4.**
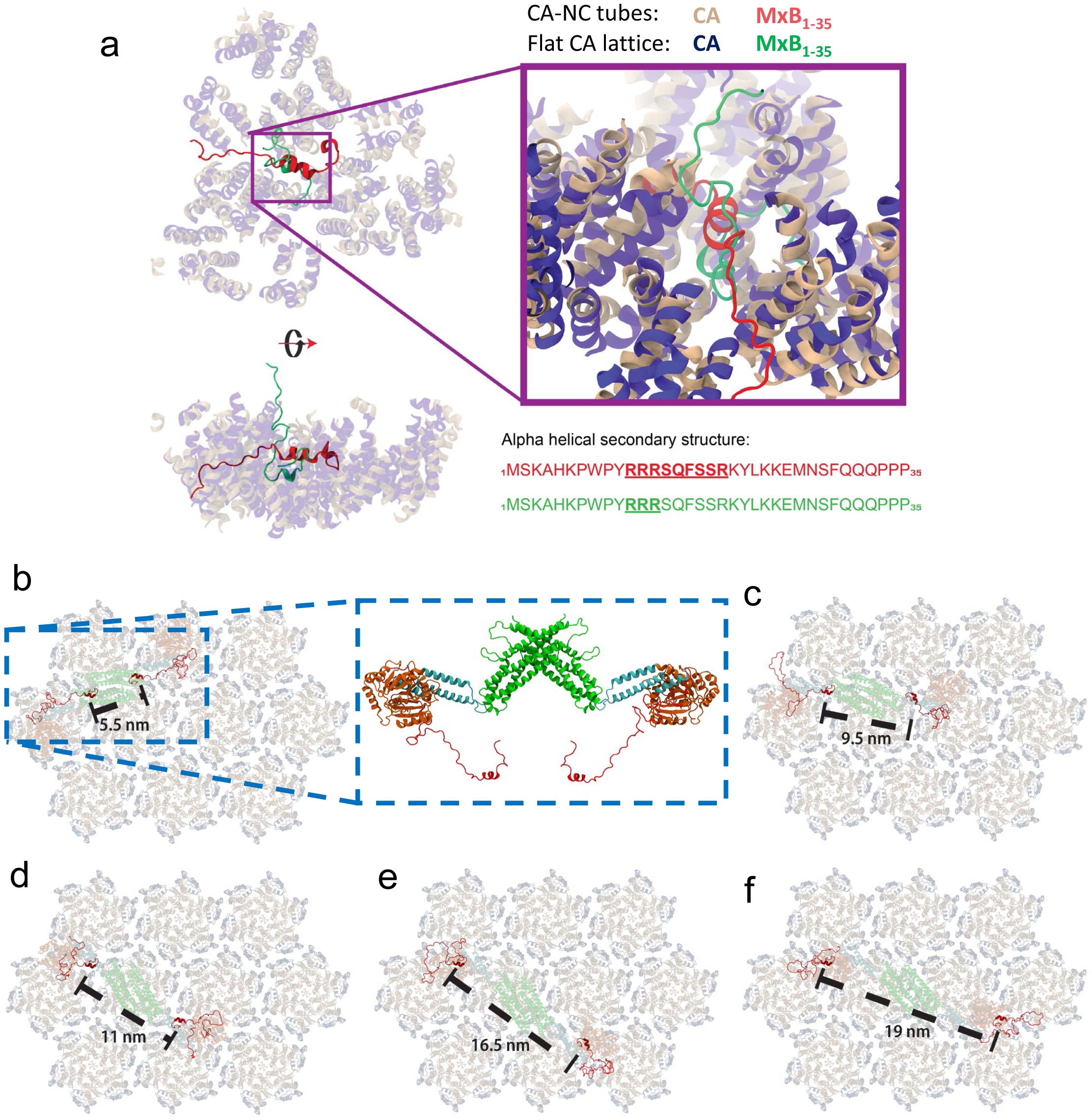
Model for MxB interaction with the curved CA lattice. (a) Comparison between MxB_1-35_ binding in curved and flat CA lattices. A representative conformation of the MxB_1-35_-CA complex from MD simulations based on the CA-NC density in presence of MxB_1-35_ (CA and MxB_1-35_ shown as ribbons in gold and red, respectively) is aligned with a representative conformation extracted from MD simulations based on a flat hexameric CA lattice (PDB ID: 4XFX) (CA in blue, MxB_1-35_ in green). In the flat lattice model, CA helices at the trimer interface are more tightly packed, and MxB_1-35_ is positioned deeper within the interface compared to its position in CA-NC tubes. During MD simulations on the curved CA lattice, MxB residues 11 to 19 adopt a well-defined α-helix, whereas in the flat CA lattice simulation, only residues 11 to 13, corresponding to the RRR motif, fold into a partial α-helix. (b-f) Illustrative models of full length MxB dimer binding modes on a CA lattice. (b) Top view of an MxB dimer bound to two adjacent CA trimer interfaces in the lattice, corresponding to a N-terminal separation of 5.5 nm. The dashed blue inset shows the full-length MxB dimer model. (c-f) MxB dimers bound to CA trimer interfaces with N-terminal separations of (c) 9.5 nm, (d) 11 nm, (e) 16.5 nm, and (f) 19 nm. These distances correspond to peaks in the pairwise distance distribution observed for MxB_1-35_ binding to CA-NC tubes. All models were derived from the crystal structure of an MxB dimer (PDBID 4WHJ) . Residues 1 to 35 were modelled based on the MxB conformations observed in MD simulations, and loops connecting the GTPase domains and the N-terminal tail domain were built using MODELLER in ChimeraX. In the CA lattice, NTD and CTD are colored gold and blue, respectively. MxB is colored by domain: N-terminal region (red), GTPase domain (orange), bundle signalling element (cyan), and stalk domain (green).

Although an MxB dimer spans approximately 20 nm between its GTPase domains (*24*), the flexible N-terminal region (residues 1–93) may allow the N-terminal binding motif to engage CA trimer interfaces separated by a range of distances. Based on our cryoET and MD results, we propose several potential binding configurations by which full-length MxB may recognize and engage the mature capsid lattice (Fig. 4b-f). The first binding configuration (Fig. 4b) corresponds to the shortest MxB pairwise distance peak (d_1_ = 5.5 nm) observed in the cryoET tomograms (Fig. 2e), in which the flexible N-terminal region enables the two MxB_1-35_ segments to bind adjacent CA trimer interfaces that share the same pair of CA hexamers. The second binding motif (Fig. 4c) corresponds to the most frequent MxB_1-35_ pairwise distance (d_2_ = 9.5 nm) measured from the cryoET fiducial analysis (Fig. 2e), with the two MxB_1-35_ segments engaging opposite trimer interfaces surrounding the same CA hexamer. The remaining proposed full-length MxB binding configurations (Fig. 4d-f) correspond to combinations of the two predominant MxB_1-35_ pairwise distances: 11 nm (2d_1_), 16.5 nm (d_1_ + d_2_), and 19nm (2d2), in which the two MxB_1-35_ segments engage trimer interfaces spanning neighbouring CA hexamers across the lattice. The proposed binding modes suggest that MxB employs a “flexible tethering” strategy, in which the N-terminal region acts as an adjustable arm that enables the dimer to sample and engage multiple CA interfaces across the capsid surface, a feature that may be critical for its lattice-sensing and restriction functions.

## Discussion

The stability, nuclear entry and uncoating of the HIV-1 capsid are tightly regulated by interactions with host factors, making the capsid a critical determinant of viral infectivity and an attractive target for therapeutic intervention (*48*). In this study, we provide a structural model of the interaction between MxB and the HIV-1 capsid, revealing how this host restriction factor exploits the unique geometry of the curved viral capsid lattice to exert antiviral activity. Based on an integrative approach combining cryoET STA with EM-guided MD simulation, we propose that the MxB N-terminus engages the capsid trimer interface through coordinated electrostatic interactions. These proposed interactions induce and stabilize an α-helical conformation in residues 10–20, representing a distinct mode of capsid recognition compared with other known restriction factors.

CryoET and STA analyses indicate that the MxB N-terminus bridges three adjacent CA hexamers at the trimer interface. Our model indicates that the conserved RRR motif forming key salt bridges with acidic residues lining this interface. This multivalent interaction explains MxB’s strong preference for assembled capsids over free CA subunits, as the complete binding surface is present only within the mature lattice. Map-back analysis further indicates that an MxB dimer can simultaneously engage two trimer interfaces, generating avidity through bivalent binding. Such crosslinking of the capsid lattice is likely to contribute to MxB’s potent restriction activity by modulating capsid stability and potentially perturbing uncoating dynamics.

The structural features of the MxB N-terminus, particularly the RRR motif, underpin its specificity and antiviral function. A similar strategy is employed by other host cofactors, such as NUP153 (*49*), which also use arginine-rich motifs to target the HIV-1 capsid. This convergence suggests that electrostatic patches at lattice interfaces represent conserved structural vulnerabilities of the capsid that can be exploited for antiviral intervention. Small molecules or peptides that mimic this binding mode may reproduce the restrictive effects of MxB, offering a potential framework for therapeutic development.

While this study provides important structural insight into how MxB engages the curved, assembled HIV-1 capsid, several functional questions remain. Future work should investigate how MxB binding influences nuclear import and reverse transcription within authentic viral cores. Live-cell imaging approaches could further elucidate the spatiotemporal dynamics of MxB-capsid interactions in their native cellular context. In addition, although our combined cryoET and computational modelling define a plausible binding mode for MxB at the CA trimer interface, the extent of the N-terminal helical conformation remains limited by the resolution of the STA density and will benefit from higher-resolution structures to refine the molecular details of the MxB-CA interface and validate the interactions proposed here. More broadly, the geometric principles of lattice recognition described here may extend to other viral systems in which capsid curvature and interface architecture govern host restriction specificity.

## Acknowledgments

We thank Dr. Yong Xiong for the MxB_1-35_-MBPdi protein. We thank Ms. Jiale Li for assistance with the cloning of CA-NC mutants. We acknowledge Diamond Light Source for access to and support of the cryo-EM facilities at the UK national electron Bio-Imaging Centre (eBIC), proposal NT29812. Computation was performed at the Diamond Light Source and Oxford Biomedical Research Computing (BMRC) facility supported by the Wellcome Trust Core Award grant number 203141/Z/16/Z with additional support from the NIHR Oxford BRC. This work was supported by US National Institutes of Health grants U54AI170791, R21AI184080; the UK Wellcome Investigator Award 206422/Z/17/Z; the UK Wellcome Discovery Award 311427/Z/24/Z; ERC AdG grant 101021133; and the Chinese Academy of Medical Sciences (CAMS) Innovation Fund for Medical Science (CIFMS), China (grant no. 2024-I2M-2-001-1); Fundamental and Interdisciplinary Disciplines Breakthrough Plan of the Ministry of Education of China (grant no. JYB2025XDXM502). This work acknowledges access to computational resources available from the SDSC Expanse supercomputer as part of ACCESS allocation MCB-170096. BioStore computational resources was made possible through funding from Delaware INBRE (P20GM103446), NIH Shared Instrumentation Grant (S10OD028725).

## Author contributions

P.Z. conceived the research. J.S. purified CA-NC protein, made CA-NC tubular assemblies, performed the co-pellet assay and prepared cryoEM/ET grids. Y.Z. and J.S. collected cryoET data. Y.Z and J.S. carried out cryoET and STA analysis. J.S.R performed density guided model building and carried out molecular dynamics (MD) simulations. J.R.P. and J.S.R analysed MD simulations. Y.Z., P.Z., J.S.R and J.R.P wrote the paper with the input from all the authors.

## Data and materials availability

All data needed to evaluate the conclusions in the paper are present in the paper and/or the Supplementary Materials. Raw tomograms and particle lists are available upon request. The cryoET STA map of CA hexamer from CA-NC tube has been deposited in EMDB under accession code EMD-54042 [https://www.ebi.ac.uk/emdb/EMD-54042]. The cryoET STA map of CA hexamer from CA-NC-MxB_1-35_ tube has been deposited in EMDB under accession code EMD-54043 [https://www.ebi.ac.uk/emdb/EMD-54043]. The cryoET STA map class1 of CA-NC-MxB_1-35_ complex has been deposited in EMDB under accession code EMD-54044 [https://www.ebi.ac.uk/emdb/EMD-54044]. The cryoET STA map class2 of CA-NC-MxB_1-35_ complex has been deposited in EMDB under accession code EMD-55015 [https://www.ebi.ac.uk/emdb/EMD-55015]. The cryoEM SPA map of CA hexamer from CA-NC-MxB_1-35_ complex has been deposited in EMDB under accession code EMD-55019 and the coordinate has been deposited in PDB under accession code 9SLY. All input parameters, scripts, coordinates and example outputs for the RosettaCM density-guided modelling and MD simulations of the CA-MxB_1-35_ complex; as well as trajectory analysis scripts are available online in Zenodo (https://doi.org/10.5281/zenodo.21252111).

## Materials and Methods

### Binding of MxB_1-35_-MBPdi to CA-NC tube

Purified MxB_1-35_-MBPdi protein was a kind gift from Dr Xiong Yong’s lab (*22*). HIV-1 CA-NC A92E was purified as previously described (*50*) in 25 mM Tris-Cl pH7.5, 150mM NaCl, 1mM ZnSO4. To assemble tubes, purified CA-NC A92E (27 mg/ml) was mixed 1:1 (v/v) with assembly buffer containing 7.5 mM Tris-Cl, pH 8.0, 150 mM NaCl and 0.4 mg/mL (TG)_50._ The mixture was adjusted to a final CA-NC protein concentration of 10 mg/mL with water and incubated overnight at 4°C. The assembled tube suspension was diluted 1:10 in buffer containing 50 mM Tris-Cl pH 8.0 and 50 mM NaCl. MxB_1-35_-MBPdi protein was added to the diluted tubes at a 1:1 molar ratio and incubated at room temperature for 1 hour.

### Co-pellet assays

MxB_1-35_-MBPdi protein was binding to assembled CA-NC A92E tube as described above at different molar ratios. After incubation, the suspension was centrifuged at 21,000 × *g* for 30 minutes at room temperature. CA-NC A92E tubes only or MxB1-35-MBPdi protein only were incubated with dilution buffer in parallel under identical conditions as control samples. Following centrifugation, the supernatant and pellet were separated and analysed by SDS-PAGE separately. Normalized volumes of the fractions were loaded to allow comparison between different binding ratios.

### CryoET sample preparation

Lacey carbon-coated copper grids (300 mesh, Agar Scientific) were glow-discharged for 45 seconds before plunge-freezing. MxB_1-35_-MBPdi bound to CA-NC A92E tubes at a 1:1 mol ratio were prepared as described above and further diluted 1:3 with 50 mM Tris-Cl pH=8.0, 50 mM NaCl followed by the addition of 6-nm gold fiducial beads.

Three microliters of the sample were applied to the carbon side of the grid, and the grid was blotted for 3.5 seconds with a blotting force of −15 before plunge freezing into liquid ethane using a Vitrobot (Thermo Fisher Scientific). The humidity was set to 100% and temperature was set to 4 °C during blotting.

### CryoET data collection

For CA-NC-MxB_1-35_-MBPdi dataset, cryoET tilt series were acquired using a Thermo Fisher Titan Krios operated at 300 keV equipped with a Gatan Quantum post-column energy filter (Gatan Inc) operated in zero-loss mode with 20 eV slit width, and Gata K3 direct electron detector. 53 tilt series were collected with SerialEM(*51*) with a nominal magnification of 64K and a physical pixel size of 1.34 Å per pixel. They were acquired using dose-symmetric range of ± 60°(*52*). The accumulated dose of each tilt series was around 123 e^-^/Å^2^ with a defocus range between −2.0 and −5.5 µm. Ten frames were saved in each raw tilt image.

For CA-NC only dataset, cryoET tilt series were acquired using a Thermo Fisher Titan Krios operated at 300 keV equipped with a Falcon 4i detector with SelectrisX operated in zero-loss mode with 10 eV slit width. 56 tilt series were collected with TOMO5 with a nominal magnification of 81K and a physical pixel size of 1.501 Å per pixel. They were acquired using dose-symmetric range of ± 60° (*52*). The accumulated dose of each tilt series was around 123 e^-^/Å^2^ with a defocus range between −2.0 and −5.5 µm. Ten frames were saved in each raw tilt image. Details of data collection parameters are listed in Table S1.

### CryoET data processing and subtomogram averaging

The automated cryoET pipeline developed in-house was used for initial tomograms (https://github.com/ffyr2w/cet_toolbox) through performing motion correction (*53*) of the raw frames, tilt-series alignment, and final reconstruction with IMOD (*54*). The fiducial markers were manually inspected to ensure the centre of predicted markers for each tilt series in eTOMO.

Initial subtomogram averaging was performed following the workflow of emClarity (*55, 56*). The CA hexamer density map (EMD-12452) (*13*) was low-passed to 30 Å and used as the initial template for template search in C1 symmetry with 6x binned tomograms with a pixel size of 8.04 Å for CANC-MxB dataset and a pixel size of 9.006 Å for CA-NC dataset. 129,046 subtomograms were selected from 53 tilt series for CANC-MxB_1-35_-MBPdi dataset and 92,814 subtomograms were selected from 56 tilt series for CANC dataset.

We converted the coordinates from emClarity to Relion-4.0 (*57*) for Relion 3D refinement in bin4 (5.36 Å pixel size for CA-NC-MxB_1-35_-MBPdi dataset and 6.004 Å pixel size for CA-NC dataset), bin3 (4.02 Å pixel size for CA-NC-MxB_1-35_-MBPdi dataset and 4.503 Å pixel size for CANC dataset) and bin2 (2.68 Å pixel size for CA-NC-MxB_1-35_-MBPdi dataset and 3.002 Å pixel size for CA-NC dataset) with C1 symmetry. Both final density maps were reconstructed at bin1 using relion_reconstruction in Relion-4.0 (*57*) and sharpened with a b-factor of −50.

For CA-NC-MxB_1-35_-MBPdi dataset, we went back to bin3 and shifted the particle coordinates from hexamer centre to the centre of tri-hexamer interface and conducted focused classification in Relion 3D classification without performing image alignment and with a mask including the central part of the tri-hexamer interface. One class with 21,392 particles with strong extra density was selected for further alignment in bin2, and final reconstruction in bin1 with a sharping b-factor of −50.

### CryoEM data collection and processing

CryoEM movie data were collected using EPU software on a Gatan K3 direct detector camera in super-resolution mode. Each movie contains 40 frames with an accumulated dose of 40 electrons/Å^2^. The calibrated physical pixel size and the super-resolution pixel size were 1.34 and 0.67 Å per pixel, respectively. The defocus was prescribed in the range from −0.8 to −2.5 μm. A total of 19,785 movies in super-resolution mode were collected for data analysis.

All frames of the raw movies were corrected for their gain using a gain reference recorded within 3 days of the acquired movie to generate a single micrograph using CryoSPARC Patch Motion Correction. The micrographs were used for the determination of the actual defocus using CryoSPARC Patch CTF Estimation. 11,355,764 particles were picked and extracted using CryoSPARC Template Picker, in which the templates were averaged from manually picked ∼1000 particles. 2,845,984 particles were selected after several rounds of CryoSPARC 2D classification. Non-uniform refinement of the final map with a global resolution of 2.77 Å was obtained with C6 symmetry applied and 3.62 Å with C1 symmetry.

### MxB_1-35_ modelling

Prior to molecular dynamics simulations of the MxB_1-35_ -CA complex, the MxB_1-35_ N-terminal tail was modelled into the cryoET/STA density using the following procedure. First, a de novo structural model of MxB_1-35_ was generated from its amino acid sequence using AlphaFold2 (*58*). Structure prediction was performed in ColabFold (*59*) using the PDB70 database for template searches, MMseqs2_uniref for multiple sequence alignment, and the alphafold_ptm weights for monomer structure prediction. Five models were generated, and the top-ranked model based on predicted local distance difference test (pLDDT) was selected for rigid body fitting into the additional density observed at the CA trimer interface in cryoET/STA maps derived from CA-NC tubes incubated in presence of MxB_1-35_-MBPdi (Fig. S6a). The docking orientation of the AlphaFold2 MxB_1-35_ model was chosen to avoid steric overlap between the MxB_1-35_ C-terminal region and the surrounding density corresponding to CA residues. The docked AlphaFold2 model served only as an initial template for subsequent density-guided model refinement. To build a model for the CA trimer-of-dimers surrounding MxB in the density, three CA hexamers were fitted into the density, and the coordinates of the CA dimers surrounding the trimer interface were extracted. Coordinates of the CA trimer of dimers were further refined via molecular dynamics flexible fitting (MDFF)(*60*) as described in previous protocols(*61, 62*). The coordinates of the docked AlphaFold2 MxB_1-35_ model were then combined with the MDFF-refined CA trimer-of-dimers model to generate a preliminary MxB_1-35_-CA complex model in preparation for RosettaCM density-guided refinement.

Only atomic coordinates of the high-confidence (high pLDDT) portion of the docked AlphaFold2 model that was well supported by the experimental density (MxB residues Arg11 to Ser18; Fig. S6a in red) were used as a template for EM density-informed comparative modelling in RosettaCM (*63*). Structure refinement of the MxB_1-35_-CA complex was performed using the Hybridizer mover in three stages. In the first stage, de novo structures for the residues not found in the template were generated and optimized in torsional and cartesian space via Monte Carlo sampling. To reduce template bias and expand conformational sampling, a residue shift of up to 4 positions was utilized, randomly varying both the sequence register and template residue coordinates during model generation . In the second stage, generated residues were connected via backbone geometry and hydrogen bonding optimization. In the third stage, sidechains were minimized using the Rosetta all-atom energy function (*64*). Throughout all stages, the atom_pair constraint scoring term was applied with weight of 1, and residue placement was guided by the STA density via the elec_dens_fast scoring term with weights of 10, 10 and 35 for stages one through three, respectively.CA residues were fixed in the first two stages of optimization to constraint the placement of newly generated MxB_1-35_ residues and to prevent MxB_1-35_ atoms from occupying CA density. During final minimization stage, CA atoms were allowed to move, enabling a broader conformational ensemble.

In total, an ensemble of 2,500 independent models was generated via this procedure (Fig. S6b). Models in the RosettaCM ensemble encompassed a diverse population of conformations, with backbone RMSDs ranging from 7.4 to 49.8 Å relative to the initial AlphaFold2 template, and a mean pairwise backbone RMSD of 14.0 ± 6.5 Å for MxB_1-35_ conformations within the ensemble (Fig. S6c). All models were subsequently minimized using the Rosetta FastRelax mover, first in torsional and then in cartesian coordinate space, with the elec_dens_fast density scoring term weighted at 35. This procedure yielded a Rosetta all-atom energy score for each structure (*64*). The minimized models were additionally compared to the STA density map via model-to-map fourier shell correlation (FSC) (*65*). The model exhibiting the highest FSC and lowest Rosetta energy score was selected as the starting structure for MD simulations (Fig. S6d-e).

### Molecular dynamics simulations

The optimized RosettaCM structure for the MxB_1-35_-CA complex was prepared for MD simulations by first placing Na+ and Cl-ions nearby the structure in places of minimum energy according to the MxB_1-35_-CA complex’s electrostatic potential via the Cionize plugin in VMD (*66*). The resulting system was then solvated in a box of TIP3P water molecules (*67*) with padding of 20 Å in each direction from the MxB_1-35_-CA complex using the solvate plugin in VMD (*66*). Finally, Na+ and Cl-ions were added to the system to achieve charge neutrality and a 150mM NaCl concentration via the autoionize plugin in VMD (*66*). The prepared simulation system encompassed ∼310,000 atoms and a total system size of ∼161 Å x 160 Å x 126 Å (Fig. S6f; Table S4).

The prepared system was then subjected to the following simulation protocol: First, energy minimization was performed on the ions and solvent while applying 100 kcal/mol harmonic restraints on protein backbone atoms to fix their positions, minimization was performed using a conjugate gradient scheme until the gradient converged to values below 10 kcal/mol⸱Å. Next, the solvent atoms were thermalized by gradually increasing the system temperature at a rate of 20 K/ns from 50 K to 310 K while restraining the positions of protein atoms as before. Once the solvent was minimized and thermalized, the harmonic spring constant for the constraints on the protein backbone atoms was reduced to 10 kcal/mol, and the minimization and thermalization steps were repeated. The constraints on protein atoms were then gradually released in a MD simulation in the NVT (constant number of particles, volume and temperature) ensemble at 310 K, in which the spring constant of the harmonic constraints was reduced at a rate of 1 kcal/mol⸱Å every 0.2 ns until the protein was completely released of constraints. As a final step, the system was equilibrated in the NPT (constant number of particles, pressure and temperature) ensemble with T=310 K and P=1 atm. After 15 ns of equilibration, the NPT simulations were extended into production runs with the same temperature and pressure. Equilibration and production runs were run in 4 independent replicates, each production run extended over 400 ns with convergence evaluated via RMSD, for a total cumulative sampling of 1.6 μs (Table S4; Fig. S8).

All simulations used a timestep of 2 fs/step and periodic boundary conditions. Long range electrostatic interactions were calculated using the particle mesh Ewald (PME) (*68*) algorithm with grid spacing of 1 Å. The cutoff for short range electrostatics and Van der Waals interactions, and the switching distance to use smoothing functions were set to 12 Å and 10 Å, respectively. All simulations used the CHARMM36m (*69*) force field for proteins, the Roux force field parameters for ions (*69*) and the TIP3P model for water molecules (*67*). Temperature control was implemented via a Langevin thermostat with a damping constant of 2 ps^-1^ , and pressure control was implemented via the Nose-Hoover barostat with a period of 100ps and decay time of 50ps. Minimization runs were performed using NAMD2.14 (*70*) while all other simulations were performed using NAMD3 alpha 13 on the GPU resident mode (*70*).

Protein-protein residue contact occupancies were calculated throughout the MD trajectories using in-house Tcl scripts implemented in VMD (*66*). Contacts between MxB_1-35_ and CA residues were defined using a minimum heavy-atom distance threshold of 3.5 Å. For each residue pair, the contact occupancy was defined as the fraction of simulation frames in which the contact was present. Contact occupancies were measured independently for each simulation replica and are reported as the cumulative occupancy over the four replicas.

### MM/GBSA binding energy calculations

The relative energetic contributions of individual MxB and CA residues to MxB_1-35_ binding were estimated by using the Molecular Mechanics Generalized Born Surface Area (MM/GBSA) method as implemented in the MMPBSA.py (*71*) module of Amber22 (*72*). MM/GBSA calculations were performed on the MD trajectories, with snapshots extracted every 0.1 ns, and per-residue free energy decomposition was used to estimate the contribution of each MxB and CA residue involved in high-occupancy contacts at the CA trimer interface. For CA residues, energetic contributions were calculated independently for each of the three equivalent chains in the trimer-of-dimers assembly, and the average across all chains is is reported in Fig. S7. Settings for MMPBSA.py included using the generalized Born solvation model (igb = 5) and with 150 mM salt concentration, and per-residue energy decomposition.

