## supplementary figures and tables for "MxB N-Terminus Adopts a Stable *α*-Helix to Engage the HIV-1 Capsid Trimer Interface"

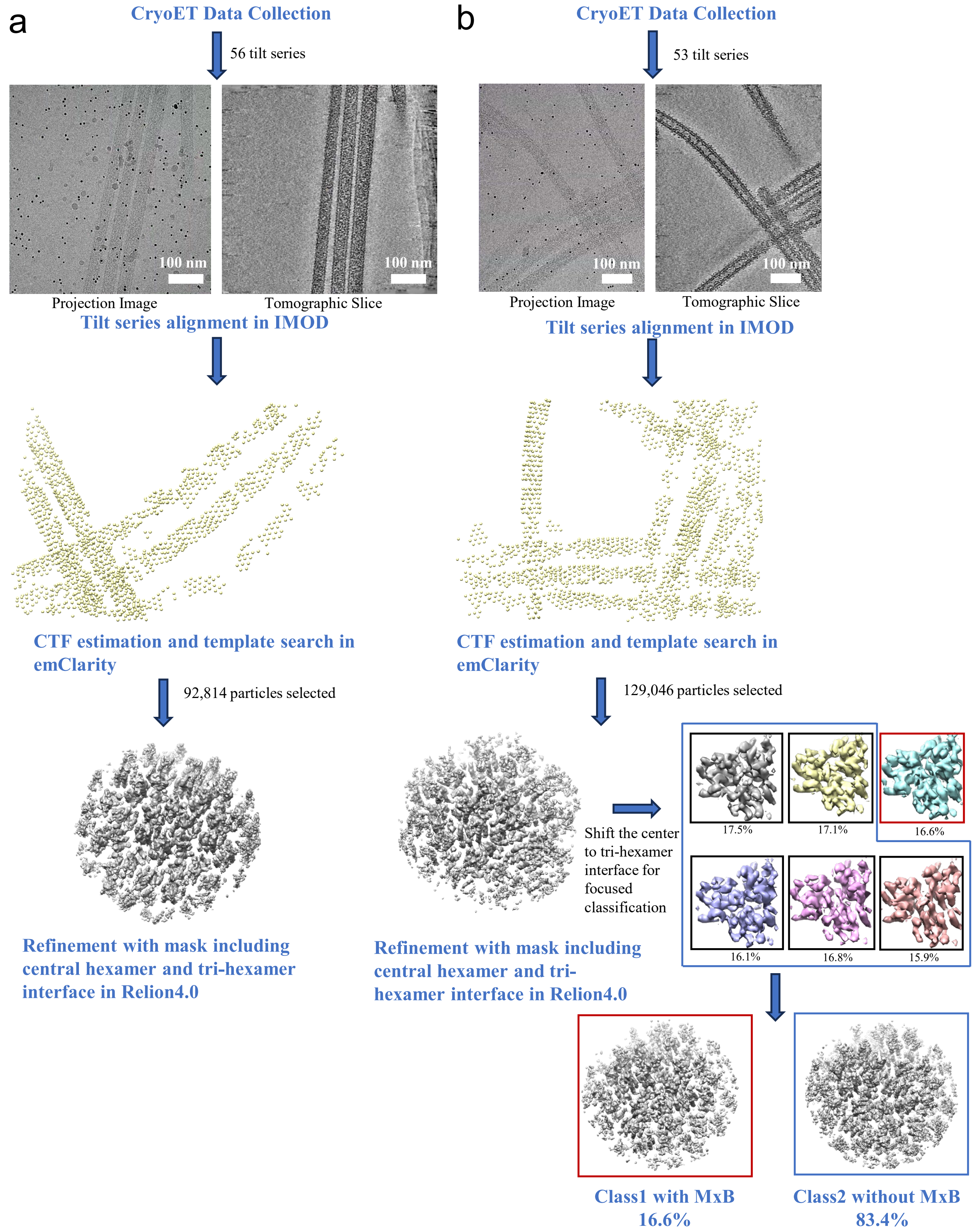


**Supplementary Figure 1 | CryoET STA data processing workflow.** (a) workflow for the CA-NC tubes. (b) workflow for the CA-NC/$MxB_{1\text{-}35}$ complexes.

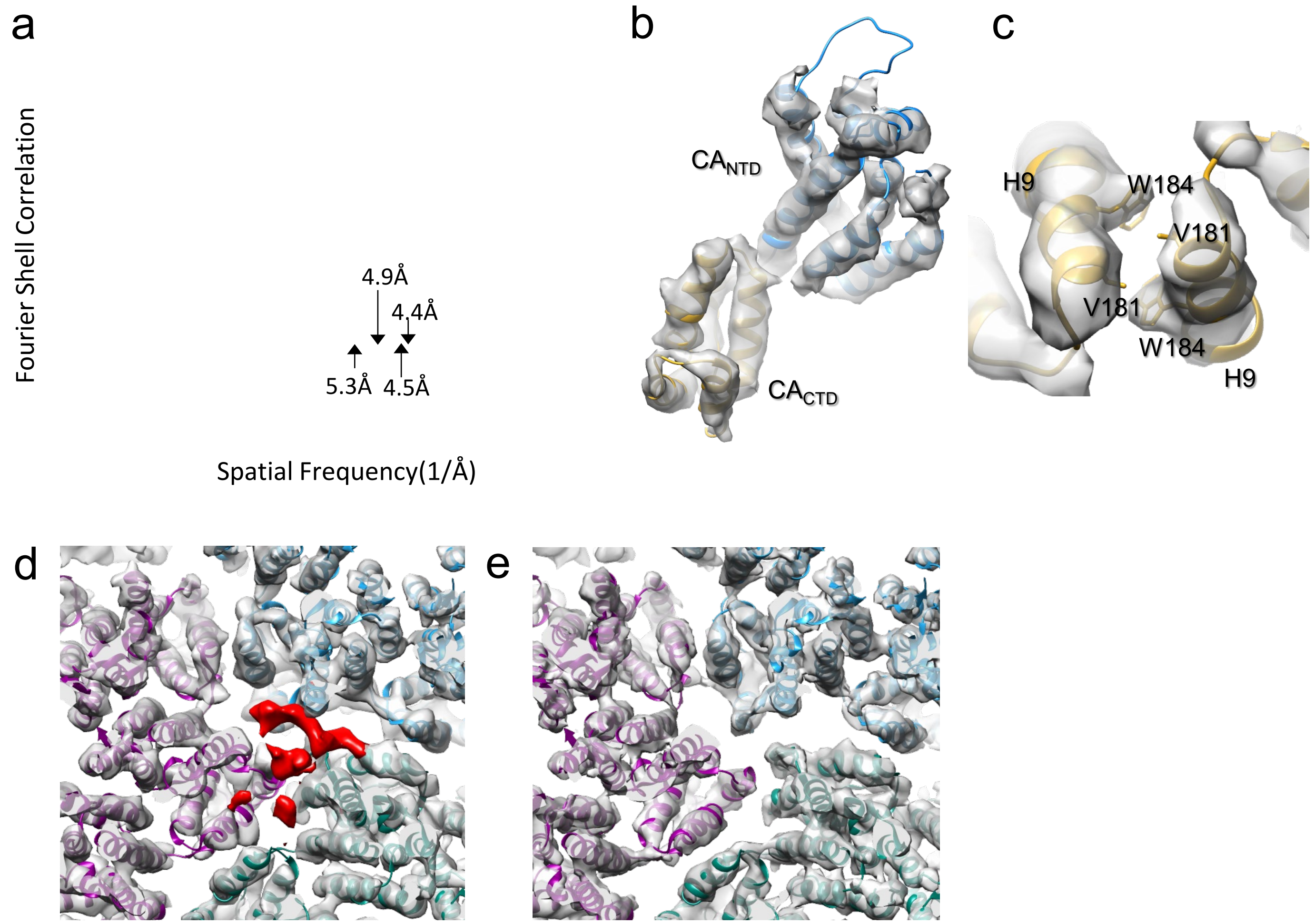


**Supplementary Figure 2 | CryoET STA analysis of $MxB_{1-35}$ and CA-NC tube complexes. Fourier Shell Correlation (FSC) plot.** (a) Fourier Shell Correlation (FSC) plot of cryoET STA density maps from CA-NC tubes (blue), CA-NC tubes in complex with $MxB_{1-35}$ (grey) and a subtomogram class with $MxB_{1-35}$ bound to the CA trimer interface (red). The resolution is indicated at the FSC value of 0.143. (b) The density map of one CA monomer from CA-NC tubes in complex with $MxB_{1-35}$ superimposed with the atomic model (PDB:4XFX) with $CA_{NTD}$ (blue) and $CA_{CTD}$ (orange) fitted separately. (c) A close-up view of the CA dimer interface from CA-NC tube in complex with $MxB_{1-35}$ , displaying the interaction between W184 and V181. (d-e) Comparison of trimer interfaces of two 3D classes from $MxB_{1-35}$ and CA-NC tube complexes superimposed with the model (PDB 9SLY). Class1 (d) distinguishes from Class2 (e) by additional densities (segmented in red). Both maps were presented at similar contour level.

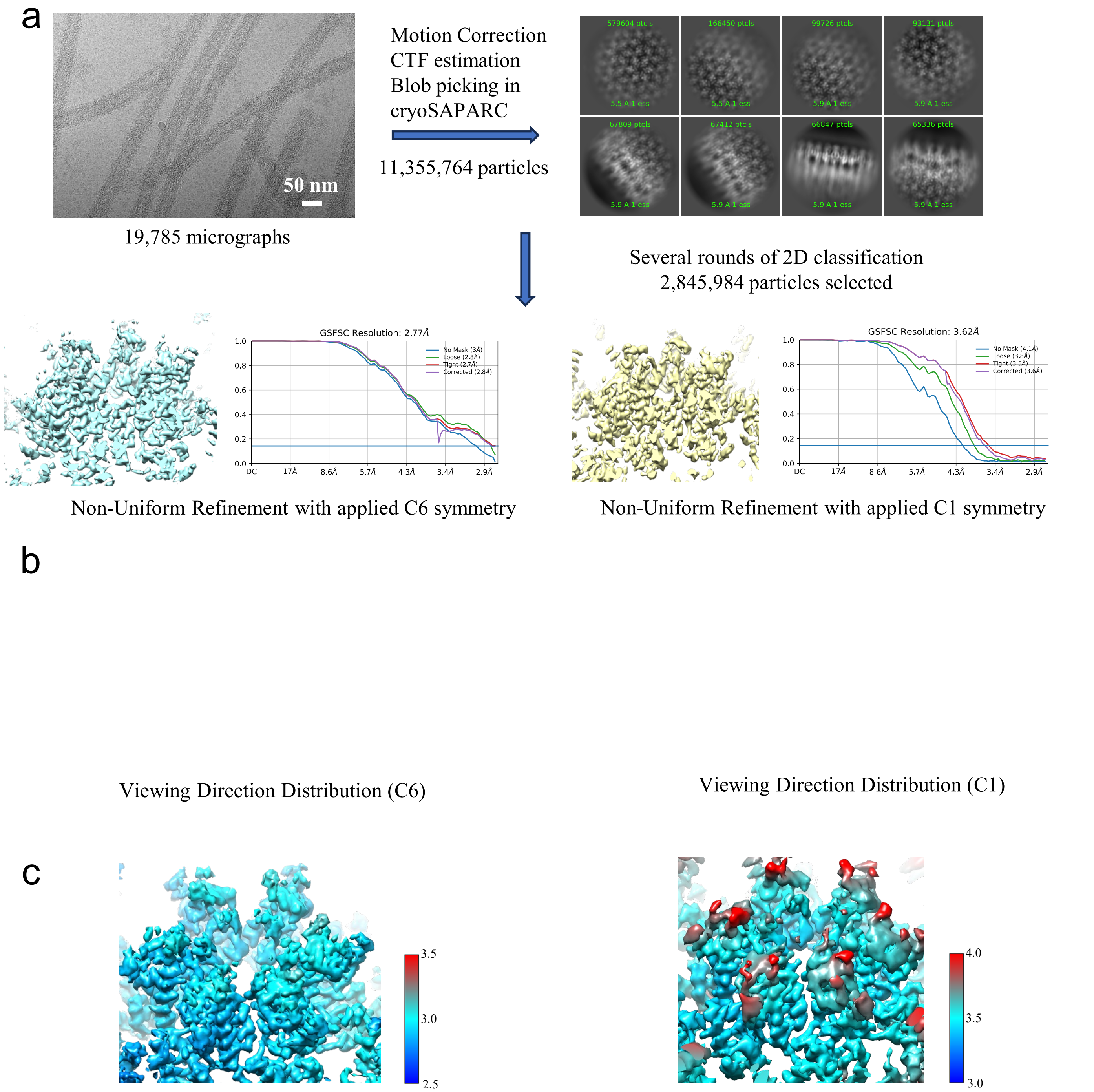


**Supplementary Figure 3 | CryoEM SPA data processing workflow.** (a) workflow for SPA data processing. (b) View direction distribution with applied C6 symmetry (left) and C1 symmetry (right). (c) Local resolution estimation with applied C6 symmetry (left) and C1 symmetry (right).

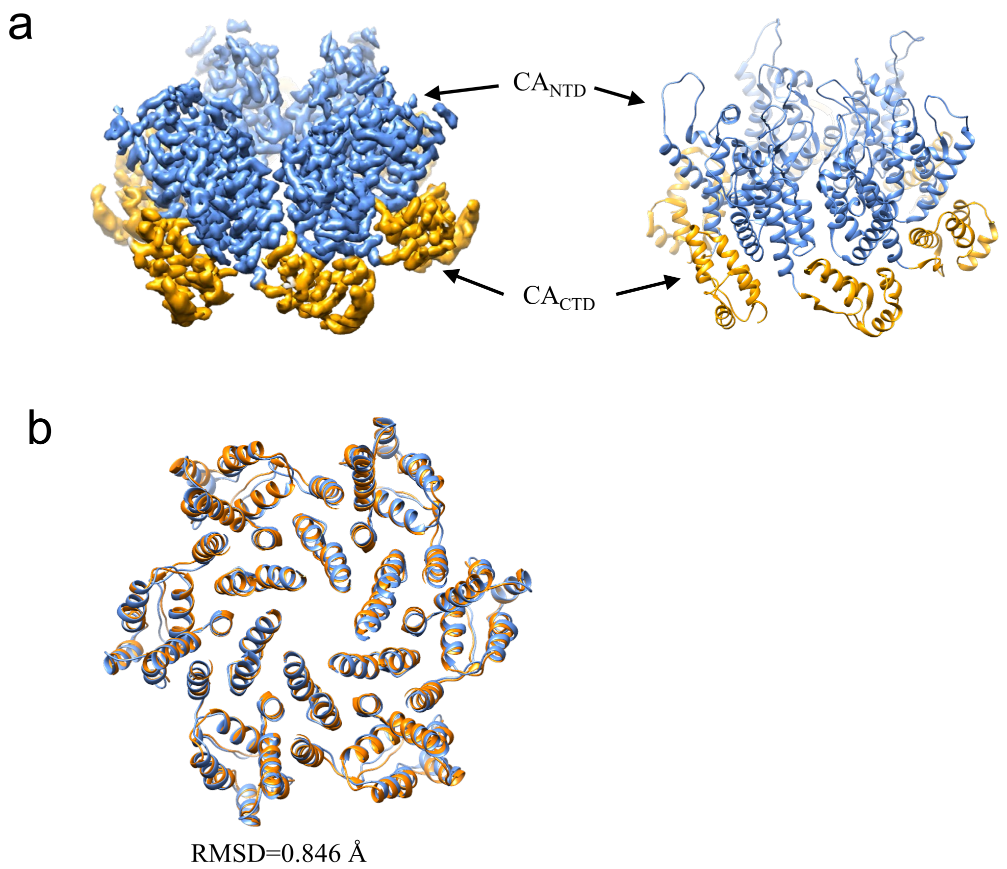


**Supplementary Figure 4 | CryoEM SPA structure of CA hexamer in MxB$_{1-35}$-bound CA-NC tubes.** (a) CryoEM map (left) and model (right) of CA hexamer from MxB$_{1-35}$-bound CA-NC tubes with CA NTD colored blue and CA CTD colored yellow. (b) Comparison of CA hexamer from MxB$_{1-35}$-bound CA-NC tubes by SPA (orange) and STA (blue).

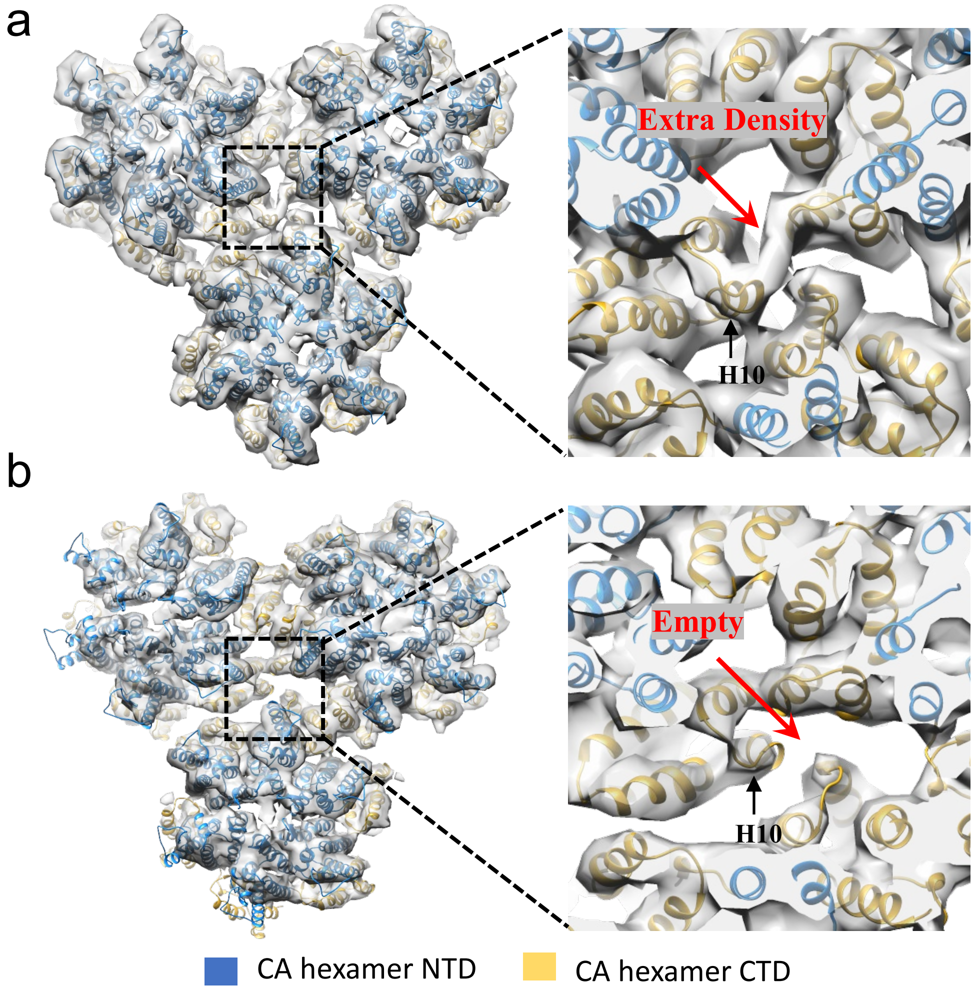


**Supplementary Figure 5 | CryoET STA density maps of CA-NC tubes and in complex with MxB$_{1\text{-}35}$.** (a) Density map of CANC tube with MxB$_{1\text{-}35}$ and extra density shown on the trimer interface. (b) Density map of CANC tube without MxB$_{1\text{-}35}$.

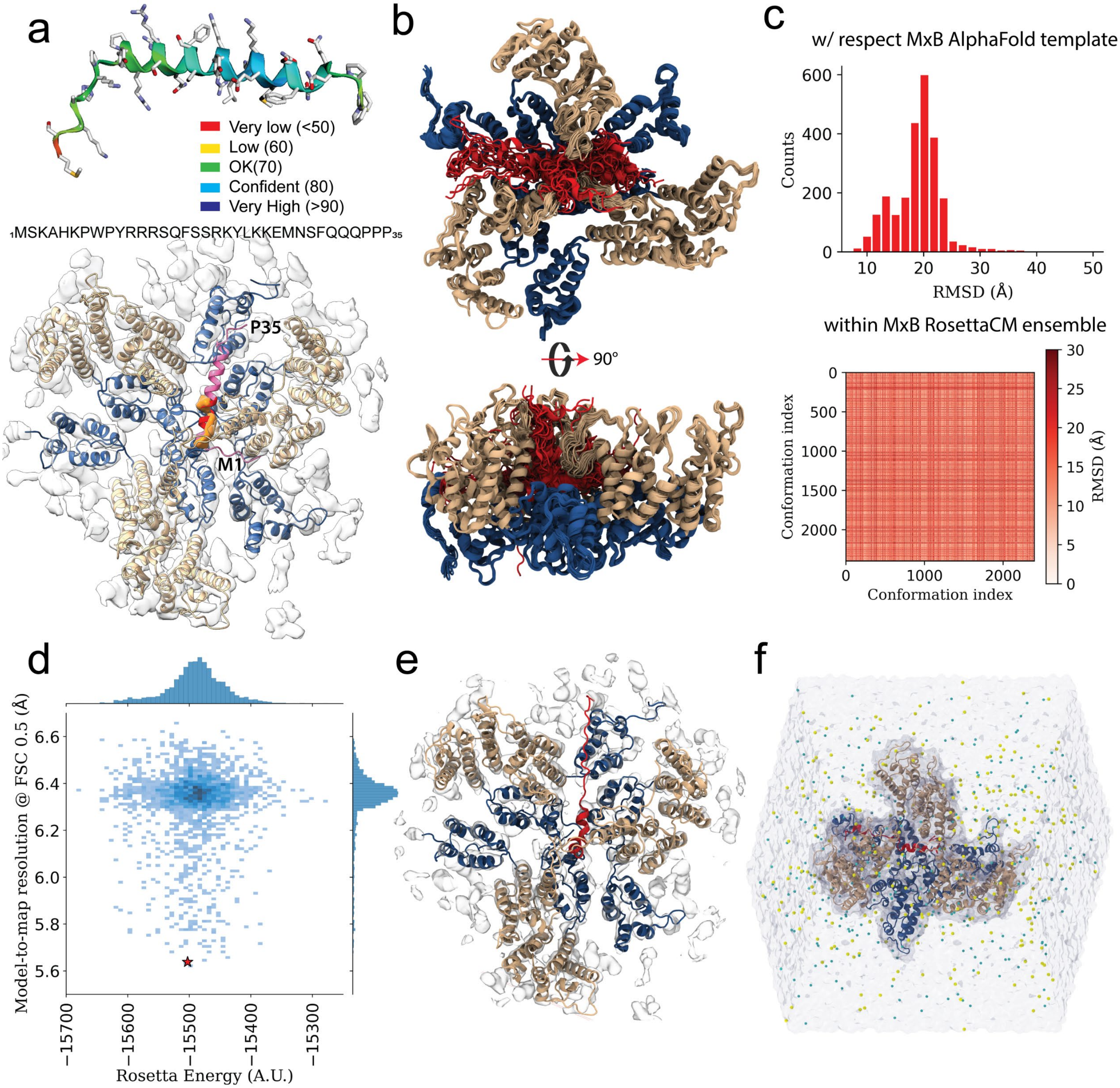

a
Very low (<50)
Low (60)
OK(70)
Confident (80)
Very High (>90)
1MSKAHKPWPYRRRSQFSSRKYLKKEMNSFQQQPPP35
P35
M1
b
90°
c
w/ respect MxB AlphaFold template
Counts
RMSD (Å)
within MxB RosettaCM ensemble
Conformation index
Conformation index
RMSD (Å)
d
Model-to-map resolution @ FSC 0.5 (Å)
Rosetta Energy (A.U.)
e
f

**Supplementary Figure 6 | Computational modeling of the $MxB_{1\text{-}35}$–CA complex.** (a) $MxB_{1\text{-}35}$ AlphaFold2 prediction colored by predicted local distance difference test (pLDDT) score (top) and the predicted structure rigid-body fitted into the STA density at the CA trimer interface (bottom). $MxB_{1\text{-}35}$, CA-NTD and CA-CTD are colored in pink, gold, and blue, respectively. MxB residues 11-18, used as a template for RosettaCM density-guided modeling, are colored in red, and corresponding STA MxB density is colored in orange.
(b) Ensemble of models generated by RosettaCM using the $MxB_{1\text{-}35}$ AlphaFold2 prediction as template. 2,500 independent models were generated; 50 representative models are shown in cartoon representation. MxB, CA-CTD and CA-CTD are colored in red, gold and blue, respectively.
(c) Backbone atom root-mean-square deviation (RMSD) of $MxB_{1\text{-}35}$ conformations generated by RosettaCM relative to the AlphaFold2 template (top) and between models within the RosettaCM ensemble (bottom).
(d) Distribution of model-to-map FSC resolution and Rosetta all-atom energy scores for the 2,500 RosettaCM $MxB_{1\text{-}35}$/CA models. The model selected as the starting structure for MD simulations, exhibiting the highest resolution and lowest Rosetta energy, is indicated with a red star in the joint plot. (e) Best scoring $MxB_{1\text{-}35}$/CA trimer-of-dimers model. (f) Ionized and solvated $MxB_{1\text{-}35}$/CA complex used for MD simulations. $MxB_{1\text{-}35}$, CA-NTD and CA-CTD are colored in red, gold and blue, respectively. Sodium and chloride ions are colored in yellow and cyan, respectively, water molecules are represented as a transparent blue molecular surface.

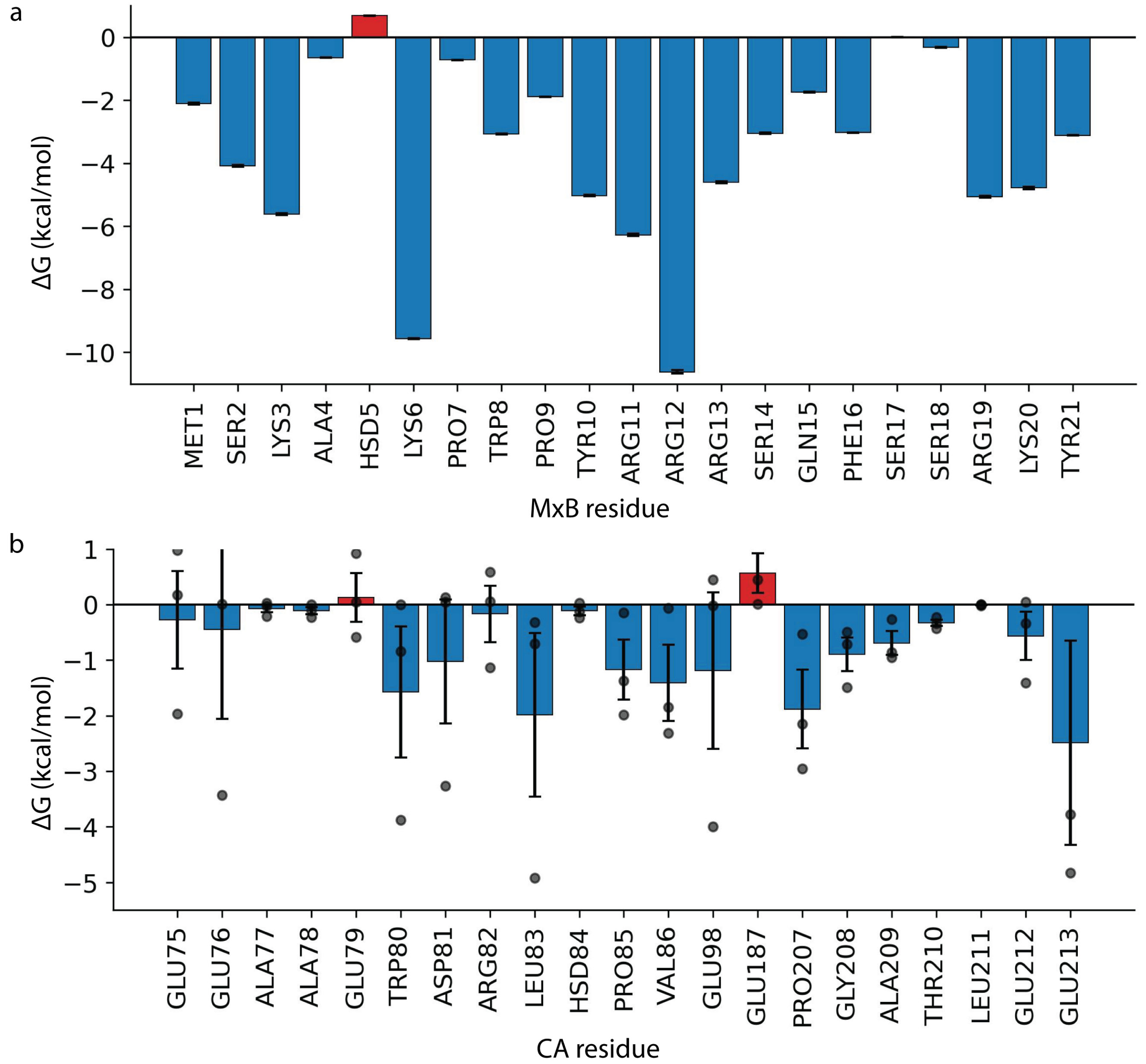


**Supplementary Figure 7 | MM/GBSA-derived per-residue energetic contributions to $MxB_{1\text{-}35}$ binding.** (a) Energetic contribution of $MxB_{1\text{-}35}$ residues that interact with CA during the MD simulations. Bar heights represent the mean binding free-energy contribution, and error bars indicate the SEM. Favorable energetic contributions (ΔG<0) are colored in blue while unfavorable energetic contributions (ΔG>0) are shown in red. (b) Energetic contributions of CA residues that interact with MxB during the MD simulations. For each CA residue, the bar height represents the average binding free-energy contribution across the three equivalent CA chains in the trimer-of-dimer assembly, and error bars indicate the standard error of the mean (SEM). Individual datapoints for each chain are shown as black circles. Favorable energetic contributions (ΔG<0) are colored in blue, while unfavorable energetic contributions (ΔG>0) are shown in red.

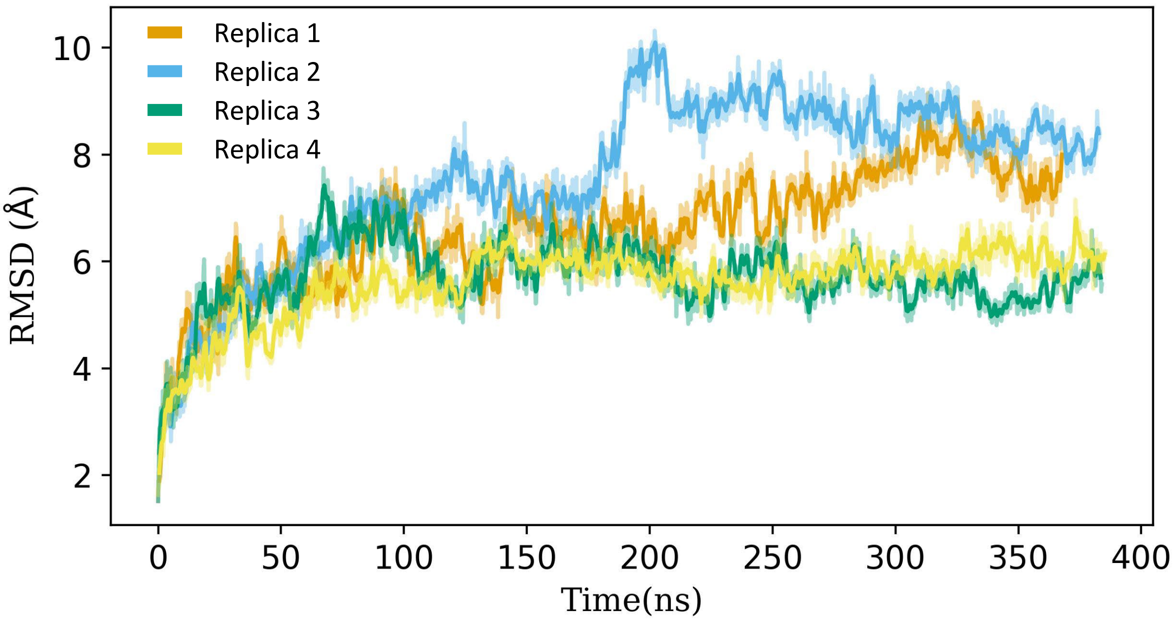


**Supplementary Figure 8 | Root-mean-square deviation (RMSD) of the CA- $MxB_{1\text{-}35}$ complex during MD simulations.** Backbone RMSD trace of the CA trimer-of-dimers is shown for each MD simulation replica. The 1ns windowed average is shown as a solid line, and the raw RMSD trace is shown with 50% transparency.

**Supplementary Table 1 |** CryoET data collection and structure determination of CA-NC/MxB$_{1-35}$ complex

| **Data acquisition** | | | | |
|---|---|---|---|---|
| Samples | CA hexamer from CA-NC tube only | CA hexamer from CA-NC/MxB$_{1-35}$ | CA tri-hexamer Class1 of CA-NC/MxB$_{1-35}$ | CA tri-hexamer Class2 of CA-NC/MxB$_{1-35}$ |
| Microscope | Titan Krios | Titan Krios | | |
| Voltage (kV) | 300 | 300 | | |
| Detector | Falcon 4i with SelectrisX | Gatan Quantum K3 Direct Electron Detector | | |
| Energy-filter | Yes | Yes | | |
| Slit width (eV) | 10 | 20 | | |
| Super-resolution mode | No | No | | |
| Å/pixel | 1.501 | 1.34 | | |
| Defocus range (µm) | -2.5 to -6 | -2.5 to -6 | | |
| Defocus increment (µm) | 0.3 | 0.3 | | |
| Acquisition scheme | Dose-Symmetric, -60/60, 3° step, group 3 | Dose-Symmetric, -60/60, 3° step, group 3 | | |
| Total Dose (electrons/Å$^2$) | 123 | 123 | | |
| Number of Frames | 10 | 10 | | |
| Number of Tomograms | 56 | 53 | | |
| **Structure determination** | | | | |
| No. of subotomograms | 92,814 | 129,046 | 21,392 | 107,654 |
| Resolution (Å) | 4.9 | 4.4 | 5.3 | 4.5 |
| Symmetry | C1 | C1 | C1 | C1 |
| b-factor applied | -50 | -50 | -50 | -50 |
| Data deposited | EMD-54042 | EMD-54043 | EMD-54044 | EMD-55015 |

**Supplementary Table 2**. CryoEM data collection and structure determination CA hexamer from $MxB_{1-35}$-bound-CA-NC tubes.

| **Data collection and processing** | |
|---|---|
| Microscope | Titan Krios |
| Magnification | 64,000 |
| Voltage | 300kV |
| Electron dose | $40e^-/Å^2$ |
| Detector | Gatan Quantum K3 |
| Defocus Range | -0.8 to -2.5 µm |
| Pixel Size | 1.34 Å |
| Symmetry | C6 |
| Particle numbers | 2,845,984 |
| Map resolution(Å) | 2.77 |
| FSC threshold | 0.143 |
| Local resolution range(Å) | 2.5-3.5 |
| **Refinement** | |
| Initial Model used (PDB code) | PDB:9I8I |
| Model resolution(Å) | 3.45 |
| FSC threshold | 0.5 |
| Map sharpening *B* factor($Å^2$ ) | -120 |
| Model composition | |
| Non-hydrogen atoms | 10350 |
| Residues | 1326 |
| R.m.s.d deviations | |
| Bond length(Å) | 0.004 |
| Bond angles(°) | 0.984 |
| Validation | |
| MolProbity score | 2.31 |
| Clashscore | 20.52 |
| Ramachandran Plot | |
| Favored(%) | 94.98 |
| Allowed(%) | 5.02 |
| Outliers(%) | 0.00 |
| Data deposition | EMD-55019 PDB:9SLY |

**Supplementary Table 3 |** Top 3 $MxB_{1-35}$ and CA contacts per residue by occupancy throughout 1.6μs MD simulation

| No. | MxB residue | Top 1 contact | Top 2 contact | Top 3 contact |
|---|---|---|---|---|
| 1 | MET1 | GLU98 (66.60%) | HIS120 (49.57%) | ILE124 (33.45%) |
| 2 | SER2 | PRO85 (68.76%) | GLU98 (62.96%) | MET96 (28.80%) |
| 3 | LYS3 | TRP80 (99.68%) | LEU83 (98.96%) | GLU98 (95.15%) |
| 4 | ALA4 | LEU83 (51.10%) | VAL86 (43.31%) | PRO85 (32.89%) |
| 5 | HIS5 | VAL86 (46.00%) | LEU83 (41.25%) | PRO85 (14.68%) |
| 6 | LYS6 | GLU79 (99.68%) | GLU76 (98.46%) | LEU83 (84.79%) |
| 7 | PRO7 | GLY208 (24.34%) | PRO207 (24.32%) | GLU187 (11.00%) |
| 8 | TRP8 | VAL86 (81.59%) | PRO85 (54.74%) | LEU83 (53.94%) |
| 9 | PRO9 | LEU83 (92.64) | VAL86 (70.64%) | GLU79 (39.39%) |
| 10 | TYR10 | ALA209 (67.60%) | GLU213 (61.89%) | LEU205 (51.39%) |
| 11 | ARG11 | PRO207 (74.15%) | THR210 (71.83%) | GLU213 (54.21%) |
| 12 | ARG12 | LEU205 (98.42%) | GLU213 (84.06%) | PRO207 (63.84%) |
| 13 | ARG13 | GLU213 (61.21%) | VAL86 (61.17%) | HIS87 (41.17%) |
| 14 | SER14 | ARG82 (84.14%) | GLU79 (61.17%) | HIS87 (41.17%) |
| 15 | GLN15 | THR210 (56.10%) | GLY208 (55.47%) | GLU75 (39.43%) |
| 16 | PHE16 | GLU213 (79.88%) | GLU212 (63.74%) | THR216 (62.06%) |
| 17 | SER17 | ARG82 (74.33%) | HIS87 (67.74%) | VAL86 (27.50%) |
| 18 | SER18 | ARG82 (94.15%) | THR110 (45.54%) | GLU75 (27.34%) |
| 19 | ARG19 | GLY208 (72.78%) | THR210 (55.00%) | GLU187 (46.97%) |
| 20 | LYS20 | GLU75 (72.34%) | GLU212 (46.47%) | LYS140 (45.74%) |
| 21 | TYR21 | GLN176 (82.71%) | ARG143 (76.08%) | LYS140 (65.93%) |
| 22 | LEU22 | THR110 (73.97%) | SER178 (65.41%) | THR108 (58.11%) |
| 23 | LYS23 | LEU136 (79.18%) | GLU76 (68.38%) | GLU79 (63.85%) |
| 24 | LYS24 | GLU113 (55.77%) | GLN179 (34.64%) | THR108 (23.51%) |
| 25 | GLU25 | GLN112 (65.63%) | LEU136 (63.53%) | GLU79 (48.21%) |
| 26 | MET26 | ARG132 (40.83%) | ILE129 (20.36%) | GLU113 (19.51%) |
| 27 | ASN27 | GLN112 (46.90%) | GLN114 (28.23%) | ARG132 (27.40%) |
| 28 | SER28 | TRP80 (25.06%) | HIS84 (23.13%) | GLN112 (12.02%) |
| 29 | PHE29 | ILE129 (60.13%) | PRO125 (54.76%) | ILE124 (53.55%) |
| 30 | GLN30 | ILE124 (24.15%) | PRO125 (15.46%) | HIS120 (11.63%) |
| 31 | GLN31 | ARG132 (37.00%) | GLU128 (21.75%) | PRO125 (20.89%) |
| 32 | GLN32 | PRO122 (8.62%) | ARG132 (8.07%) | ARG100 (7.77%) |
| 33 | PRO33 | ILE135 (25.51%) | ARG132 (20.61%) | SER41 (12.65%) |
| 34 | PRO34 | ARG132 (29.97%) | ILE135 (29.94%) | LEU136 (28.66%) |
| 35 | PRO35 | GLN179 (17.50%) | ALA88 (7.25%) | VAL86 (7.19%) |

**Supplementary Table 4 |** Summary of the CA-$MxB_{1-35}$ complex system built for MD simulations.

| System description | Dimensions (Å) | Total number of atoms | Total number of water molecules | Salt concentration (mM NaCl) | Simulation length (ns) | No. replicas |
|---|---|---|---|---|---|---|
| CA trimer-of-dimers in complex with $MxB_{1-35}$ NPT equilibration/production. | 160.4 × 163.9 × 117.7 | 312,678 | 96,637 | 150 | 400 | 4 |
